# Cryopreservation of Sea Urchin (*Lytechinus pictus*) Embryos and Development Through Metamorphosis

**DOI:** 10.1101/2025.07.23.666466

**Authors:** Victor D. Vacquier, Amro Hamdoun

## Abstract

**Background:** Sea urchins have contributed to knowledge of fertilization, embryonic development and cell physiology for 150 years. Their evolutionary position, as basal deuterostomes, and their long background in developmental biology, motivate establishing a genetically-enabled sea urchin species. Because of its relatively short generation time of 4-6 months and ease of culture, our lab has focused on the California sea urchin *Lytechinus pictus* as a multigenerational model, and produced knockout and transgenic lines using this species. To ensure that diverse genetic lines can be preserved, methods must be developed to cryopreserve gametes and embryos. We have previously reported methods for cryopreservation of sperm, but robust methods to preserve embryos remained lacking.

**Results:** Here, we describe a relatively simple method to cryopreserve late gastrulae embryos of *L. pictus*. Importantly we show that, after thawing and culturing, the embryos progress through larval development, undergo metamorphosis and yield juvenile adults, indicating the method is robust.

**Conclusion:** The cryopreservation of embryos is an important advance that will facilitate the biobanking, sharing and long-term preservation of diverse genetic lines. This method may also eventually prove useful for cryopreservation of embryos of other marine invertebrates.

## 1 INTRODUCTION

Sea urchins are basal deuterostomes, the evolutionary line leading to the vertebrates. The study of their gametes and embryos was central to the foundational knowledge of cell and developmental biology. A renewal and expansion of sea urchin research is currently underway, spurred on by the development of CRISPR gene editing and rapid gene sequencing. The renewed interest in sea urchins as experimental models includes studies of: the evolution of cell fate during development^1^, innate immunity^2-5^, drug transporters^6,7^, biomineralization^8-11^, animal longevity^12^, metamorphosis^13,14^, animal electrical oscillations^15^, and new drug discovery^16-19^.

Moving towards a genetically enabled echinoderm model has long been a goal of the field^20^. Our laboratory is focusing on the California sea urchin, *Lytechinus pictus* (Lp) as a multigenerational echinoderm model, because of its relatively short generation time of 4-6 months and ease of culture^21-23^. As genetic lines become common, robust procedures for cryopreservation and dissemination of lines will be needed.

Approximately 10 sea urchin species have had their embryos cryopreserved by various procedures and this previous work has been extensively reviewed^25-30^. One hallmark of the above referenced reviews is that cryopreservation methods for sea urchin embryos can be species-specific. And, importantly, in only a few of these studies have the thawed embryos been raised through metamorphosis to juvenile adults of unknown numbers. A robust procedure for producing juveniles is essential for propagation of urchin lines.

To preserve laboratory-constructed genetic lines for our use, and for sharing with others, we previously developed a procedure to cryopreserve Lp sperm^24^. To further expand capabilities for cryopreservation, we report here a new method to cryopreserve late stage Lp gastrulae that, when thawed, develop into feeding larvae that undergo metamorphosis to yield juvenile adult sea urchins.

## 2. RESULTS AND DISCUSSION

### 2.1. Rationale for developing the new method of embryo cryopreservation

Our goal to develop *Lp* as a new multigenerational genetic model is progressing. One future need for this project is a new cryopreservation method that would yield adequate numbers (hundreds) of individuals for banking genetic lines, sharing lines with other labs and for screening specific genes.

### 2.2. Success of cryopreservation of *Lytechinus pictus*

In the experiment result shown in Fig. 1, after thawing approximately 1545 embryos and five days of culture, 1028 four-arm feeding Lp plutei larvae were selected by hand and cultured in 1 liter FSW. After an additional 13-18 days 510 of these larvae were transferred to conditioned plates to induce metamorphosis. This produced the 461 juveniles shown in Fig. 1 at low magnification. Results of another experiment are shown at higher magnification in Fig. 2 to show normal morphology. Four additional experiments yielded similar results (Table 1).

**Figure 1.**
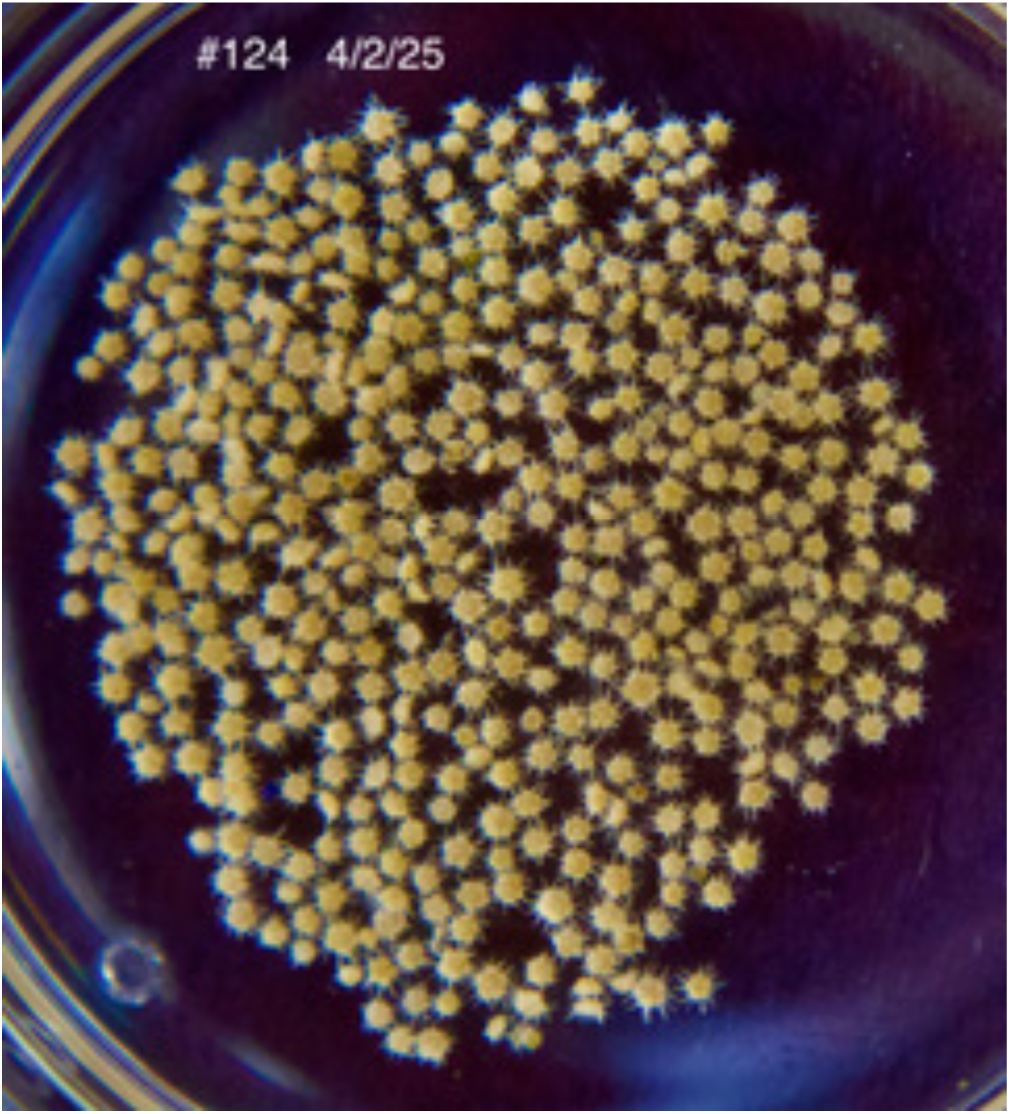
After 65 days in liquid nitrogen atmosphere 1 ml containing approximately 1545 late gastrulae was thawed and the embryos rehydrated and cultured in a 1 liter beaker with a 15 rpm stirring motor. Between 18 and 25 days of culture 510 mature larvae with large adult rudiments were removed and put on conditioned plates to induce metamorphosis. These 461 juvenile sea urchins were fixed in 2% paraformaldehyde in FSW seven days post metamorphosis.

**Figure 2.**
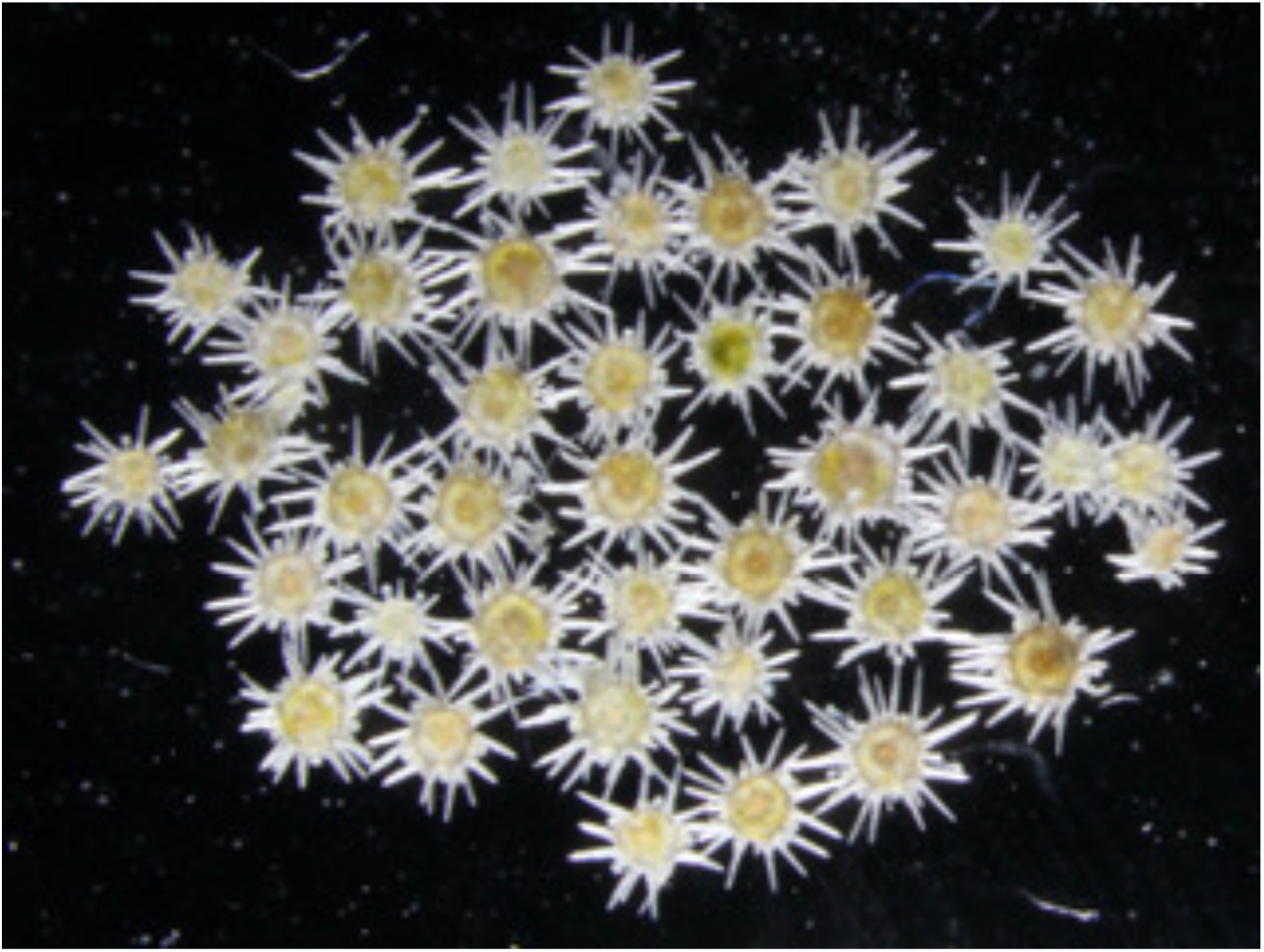
These 41 juveniles were stored in liquid nitrogen atmosphere for 200 days before thawing rehydration, culturing and metamorphosis. They are presented at higher magnification from Fig.1 to show the normal morphology of one week post metamorphosis juveniles. They are fixed in 2% paraformaldehyde in FSW.

**Table 1.** Results of five other experiments. Exp, Experiment number; dLN, days in the liquid nitrogen atmosphere; E ReHy, approximate number of late gastrulae rehydrated into FSW. Juve, number of juvenile sea urchins; % Juvie produced from the rehydrated late gastrulae.

| **Exp** | **Days in LN** | **E ReHy** | **Juvie** | **%Juvie** |
| --- | --- | --- | --- | --- |
| 127 | 77 | 1000 | 141 | 14 |
| 123 | 125 | 785 | 250 | 32 |
| 119 | 133 | 1270 | 272 | 21 |
| 94 | 323 | 750 | 194 | 26 |
| 94 | 365 | 750 | 180 | 24 |

### 2.3. Abnormalities

Within two hours after thawing, rehydration and return to 100% FSW, essentially 100% of gastrulae are motile. There appears to be a 1-2 day delay before the first pair of arms begin to bud from prism stage embryos. The larval plutei begin to feed at day 4 after return to 100% FSW. By day 13 the beginning adult rudiment appears on one side of the larval stomach^31-33^. The most frequent abnormalities induced by the freeze-thaw-rehydration are: lack of symmetry and length of the first pair of arms, failure of plutei to feed and failure to produce an adult rudiment. The large number of larvae available in the sea urchin system makes these abnormalities inconsequential to the value they provide.

### 2.4 Flexibility in formulation of solutions

In the methods section we present reagent concentrations that are optimal for cryopreservation of Lp embryos. Researchers interested in trying our cryopreservation method on other species of sea urchins, or other marine invertebrates, should know that our optimal formulation given below has flexibility. For example, in the extender solution (EXT), 85% FSW/15% 1M sucrose (v/v), can be 70% FSW/30% 1M sucrose. Ten mM HEPES can be raised to 15 mM be adjusted to pHs 6.8 to 7.4 and still be successful.

The PVP360K can be 5-10 mg/ml, but concentrations higher than 10 mg/ml do not improve results. Ten mg/ml soluble protein can be any mammalian serum albumin, or chicken ovalbumin. Lastly, in the cryopreservation solution (CPA), DMSO concentration can vary from 14% v/v (2 M) to 20% v/v (2.8 M).

## 3 EXPERIMENTAL PROCEDURES

### 3.1 Biological Material

Adult *Lytechinus pictus* (Lp) were collected at San Diego, California, kept in aquaria (22°C) exchanging with the ocean and fed on the giant kelp, *Macrocystis pyrifera*. Adult urchins were spawned by injection of 0.55M KCl and eggs fertilized and cultured at 22°C at concentrations of 1,000-2,000 zygotes in 8 ml of SW in a Petri plate of 5 cm diameter. At 23-26 hours (22-23°C) embryos are mid to late gastrulae^31-33^. Late gastrula stage, just at the beginning of the prism stage, is the optimal time to harvest the embryos. All seawater (FSW) was filtered through a 0.22 μm membrane. The following protocol produced the juveniles presented in Fig. 1.

### 3.2 Cryopreservation

Late gastrulae were collected by pouring small Petri plate cultures into a 50 ml conical tube, which was hand centrifuged gently to sediment the embryos. The supernatant was aspirated away and the embryos resuspended in fresh FSW. The number of embryos per ml was determined by streaking 100μl of embryo suspension on a dry Petri plate and counting the embryos per streak in 10 streaks and then averaging. The desired number of embryos were sedimented again, the supernatant completely aspirated away, and the embryo pellet resuspended at 1,000-3,000 embryos per ml in the extender solution (EXT). The EXT formula was: 85% FSW/15% 1 M sucrose v/v, containing 10 mM HEPES buffer pH 7.4, 10 mg/ml polyvinyl pyrrolidone (360,000 ave mol wt, Sigma, PVP) and 10 mg/ml rabbit, or bovine serum albumin, or chicken ovalbumin, and 0.2 mg/ml horse radish peroxidase. The pH was adjusted to 7.4 with 1N NaOH. One ml aliquots of embryo suspension in 100% EXT were distributed into wells of a 12 well culture plate (Corning REF3513) that was placed on an orbital shaker table set at 120 rpm. The embryos were rotated in EXT for 20 min before addition of the cryoprotective agent (CPA), which was: 84% EXT/16% v/v (2.3M) dimethyl sulfoxide (DMSO, Fisher Scientific). One addition of 100 μl CPA was added to the 1 ml of rotating wells at 2 or 3 min intervals without stopping rotation. After 10 additions of CPA the total volume in each well was 2.0 ml and the DMSO concentration was 1.15 M. After addition of the 10^th^ aliquot of 100 μl CPA the plate was rotated an additional 20 min. Two ml capacity cryovials (Corning) were labelled and 1.1 or 1.6 ml from each well was put into each cryovial and caps sealed tight using pliers to hold the cryotube base. Cryotubes were put into a cardboard 81 position cryobox and placed in the LN atmosphere of the Dewar above the LN liquid phase. No attempt was made to study the rate of cooling, or timed steps at different temperatures.

### 3.3. Thawing and rehydration

At various times after storage in the LN Dewar, cryotubes were removed and within 60 sec, completely emersed in a 50°C water bath for 20 sec, then instantly emersed in 18°C water for 4 min. The tubes were rapidly moved by gloved hand during both emersions. The thawed tubes were laid on crushed ice and within 5 min 1.0 ml of cryotube contents transferred to wells of a 12 well culture plate and rotation started at 120 rpm. The rehydration solution was 98% FSW/2% 1M Dextrose v/v with 2 mg/ml of chicken ovalbumin (or BSA), 0.2 mg/ml horse radish peroxidase and 10 mM HEPES buffer adjusted to pH 8.0 with 1N NaOH (All chemicals were from Sigma Aldrich). The rehydrating solution was added at 1 min intervals without stopping rotation, and after each fifth addition, the volume of the solution was increased. The five additions of each volume in μL were: 50, 75, 100, 150, 200, 250 and 300. This stepwise addition takes 35 min and adds 5.6 ml to the 1.0 ml of thawed embryos in each well. After the last 300 μL addition, the rotation was continued for 15 min. and then the 6.6 ml volume of each well was transferred to a 12 ml conical glass tube and 5 ml FSW added in 1 ml aliquots. The 12 ml tube was inverted 5 times after each 1 ml addition. The embryos were sedimented by gentle hand centrifugation and the supernatant removed by aspiration. Eight ml FSW was added to each tube and the tube inverted 5 times to wash the embryos. The embryos are then recentrifuged by hand, the wash FSW aspirated away, and the embryos resuspended in 10 ml FSW and poured into a standard size Petri plate (10 cm diameter) containing 20 ml FSW.

### 3.4 Culturing and metamorphosis

The experiment yielding Fig.1 provides a good example. After 2 days of culturing the Petri plate was observed with a stereo dissecting microscope and 1028 of the best looking gastrulae were removed using a 9 cm glass pipette and placed in a one liter beaker with 900 ml FSW, which was stirred with a 15 rpm TYC electric motor driving a 2.5 × 5 cm plastic paddle held 2 cm from the beaker bottom. The single cell alga *Rhodomonas lens* was added daily^32-33^. Penicillin G and Streptomycin Sulfate (Sigma Aldrich, St Louis Mo) were both added to the beaker at 50 mg each. Seven hundred ml of the FSW was changed every 3^rd^ day and fresh antibiotics added. By 18 days the larvae were close to maximum size with large adult rudiments. By 21 days some advanced larvae had undergone spontaneous metamorphosis. The signs of metamorphosis competency are: retraction of arms to 50% of length, shrinkage of the entire body, development of intense yellow color, greenish color of the stomach and movement of the tube feet of the adult rudiment^31-33^. Any two of these visual attributes defines a larva as being metamorphosis competent. Visually competent larvae were removed by hand one-by-one and placed in 10 cm diameter “conditioned” Petri plate that had been stored for weeks in unfiltered, flowing seawater and was encrusted with a thick microbial film of unknown composition. Within minutes to hours most competent larvae undergo metamorphosis. Juveniles are then moved to a clean 10 cm Petri plate and feed on the alga Nitzschia alba^31-33^ and grown to sexual maturity as described^31-33^

## Abbreviations

FSW: filtered seawater
Lp: *Lytechinus pictus*
LN: liquid nitrogen
#PVP360K: polyvinyl pyrrolidone 360,000 MW
EXT: extender solution
CPA: cryoprotective agent
DMSO: dimethyl sulfoxide.

## ACKNOWLEDGMENTS

We thank Karen Ong for her assistance with algal culturing.

